# A hit-and-run strategy for protoplast reprogramming and regeneration into transgene-free plants

**DOI:** 10.64898/2026.04.10.717603

**Authors:** Gajendra Singh Jeena, Mehrun Nisha Khanam, Chulmin Park, Bosl Noh, Yoo-Sun Noh

## Abstract

The ability of protoplasts to regenerate into whole plants underpins advances in crop engineering and pluripotency research. However, most protoplasts exhibit poor division and limited shoot regeneration, restricting their broader utility. Here, we present EppTec (**E**fficient regeneration of transgene-free whole **P**lants from **P**rotoplasts reprogrammed by **T**ransiently **E**xpressed **C**ombinatorial factors), a transient expression platform using defined combinatorial factors (CFs) to unlock protoplast pluripotency and enable transgene-free whole-plant regeneration. Among 24 CFs tested, co-transfection of SEPW (SCR, ESR1, PSK5, and WOX5) markedly enhanced regeneration across diverse plant species in an evolutionarily conserved manner. We demonstrate that SEPW co-transfection induces epigenomic reprogramming, formation of a distinct cell cluster undergoing reprogramming and cell-cycle re-entry, and long-term reprogramming into pluripotent cells. These findings establish EppTec as a robust strategy to restore the regenerative capacity of plant cells from diverse species. EppTec may serve as a platform to revolutionize regeneration-based plant biotechnologies and conserve endangered plant species.

## Introduction

One of the fundamental questions in biology is unravelling the intricate mechanisms governing the reprogramming of differentiated somatic cells to pluripotent or totipotent states. Plants exhibit remarkable reprogramming potential, enabling the regeneration of organs or entire plants from diverse sources, including tissues and individual cells (Ikeuchi et al. 2019). Over the past century, research has expanded on Haberlandt’s concept of totipotency, postulating that specialized plant cells retain the ability to differentiate into various cell types (Haberlandt et al. 1902). This principle remains central to plant research and has ample implications for agriculture and biotechnology.

Protoplasts are capable of reconstructing cell walls and undergoing division, making them invaluable tools for genetic engineering and studying cellular processes (Yokoyama et al. 2016). Mature mesophyll cells in intact plant tissues usually have limited division and regeneration capabilities. However, mesophyll protoplasts can reconstruct cell walls, reinitiate cell cycles, generate calli, and initiate regeneration processes (Takebe et al. 1971). Although Arabidopsis mesophyll protoplasts divide to form microcalli in a gel-based medium, their regeneration rate is notably low (Dovzhenko et al. 2003). A variety of protoplast regeneration protocols have been developed for many plant species, including Arabidopsis. However, challenges persist, as exemplified by low shoot-regeneration rates, revealing unique technical hurdles specific to protoplast regeneration compared to tissue explant-based regeneration (Jeong et al. 2021). Additionally, the intricate molecular mechanisms orchestrating cell-fate transitions during protoplast regeneration remain elusive. Nevertheless, protoplast-regeneration techniques are widely used in various genetic modification studies, such as gene transformation and genome editing in plants, albeit with limited success (Reed and Bargmann 2021). Therefore, the regeneration process of protoplasts stands as a pivotal bottleneck constraining the widespread deployment of genome-editing tools in plant biotechnology, which may present avenues for enhancing agricultural productivity (Altpeter et al. 2016).

Although earlier strategies employed chemical cues to promote protoplast regeneration, they were limited by a lack of specificity (Jeong et al. 2021; Ford K.G. 1990; Mathur et al. 1995; Chupeau et al. 2013; Xu et al. 2021; Sakamoto et al. 2022). In contrast, transcription factor (TF)-mediated reprogramming offers a precise and targeted approach by directly modulating key developmental pathways to drive regeneration. Recent advances have highlighted the role of specific transcription factors, such as GRF-GIF, in enhancing shoot regeneration efficiency, underscoring the potential of genetic regulators in cellular reprogramming (Luo et al. 2021). Building on this foundation, our study investigates the ability of defined factors, including TFs, to induce protoplast reprogramming and regeneration, providing a mechanistic framework for improving plant regeneration strategies. Here, we introduce EppTec (**E**fficient Regeneration of Transgene-Free Whole **P**lants from **P**rotoplasts Reprogrammed by **T**ransiently **E**xpressed **C**ombinatorial Factors), a hit-and-run strategy that robustly induces protoplast reprogramming and regeneration into transgene-free whole plants across both dicot and monocot species. EppTec markedly enhances regeneration efficiency and speed, offering broad utility for regeneration-based plant biotechnologies.

## Results

### Transient transfection enables sustained gene expression in Arabidopsis mesophyll protoplasts

In this study, we focused on the transient expression of combinatorial factors (CFs) in Arabidopsis mesophyll protoplasts, known for their full differentiation and homogeneity compared to protoplasts of other origins. Protoplasts isolated from the leaves of 15-day-old seedlings showed high viability (> 95%) as assessed by fluorescein diacetate (FDA) staining (Supplementary Fig. S1A), and remained largely viable (∼73%) after 7 days (7d) in protoplast induction medium (PIM). Despite this, only 9% of viable cells re-entered the cell cycle and formed microcalli (≥ 4 cells), with 14% undergoing a single division and 4% undergoing two divisions. A small fraction (∼3%) exhibited abnormal or arrested morphologies in PIM (Supplementary Fig. S1B). To examine early regenerative events, we tracked cell wall reconstruction and expansion using calcofluor white (CW) staining, which was detectable at day 2 (d2). The first division occurred at d4, progressing to multicellular microcalli at d7 (Supplementary Fig. S1C). Following standard regeneration protocols, microcalli formed in PIM at d11 increased in size under sequential callus induction medium 1 (CIM1) and CIM2 conditions until d71 and ultimately resulted in the regeneration of whole plants at d120 through shoot induction medium (SIM) and root induction medium (RIM) cultures (Supplementary Fig. S1D). We also evaluated two shortened regeneration protocols to accelerate the protoplast culture timeline (Supplementary Fig. S1, E and F).

To assess transgene expression dynamics, we transfected protoplasts with HBT-PGSG-NOS plasmids expressing sGFP, mRFP, or mBFP fluorescence markers. These plasmids lack plant genomic integration elements and plant replication origins, but are replication-competent in *Escherichia coli*. Fluorescence was first detectable at d1 and persisted by d7 post-transfection, with ∼90% of cells showing a strong signal at d1 (Fig. 1, A and B). Signal intensity remained stable through d7, declining by > 60% at d9 and becoming undetectable by d11 (Fig. 1C). These results demonstrate the high transfection efficiency and transient expression stability of our transfection system. Given that pluripotency-conferring cell reprogramming in plant tissue culture typically establishes within 4∼14d, the 7d expression window might offer a feasible platform for cell reprogramming. We also observed similar expression dynamics for the designed factor-expressing constructs (Supplementary Fig. S2, A to F).

**Figure 1.**
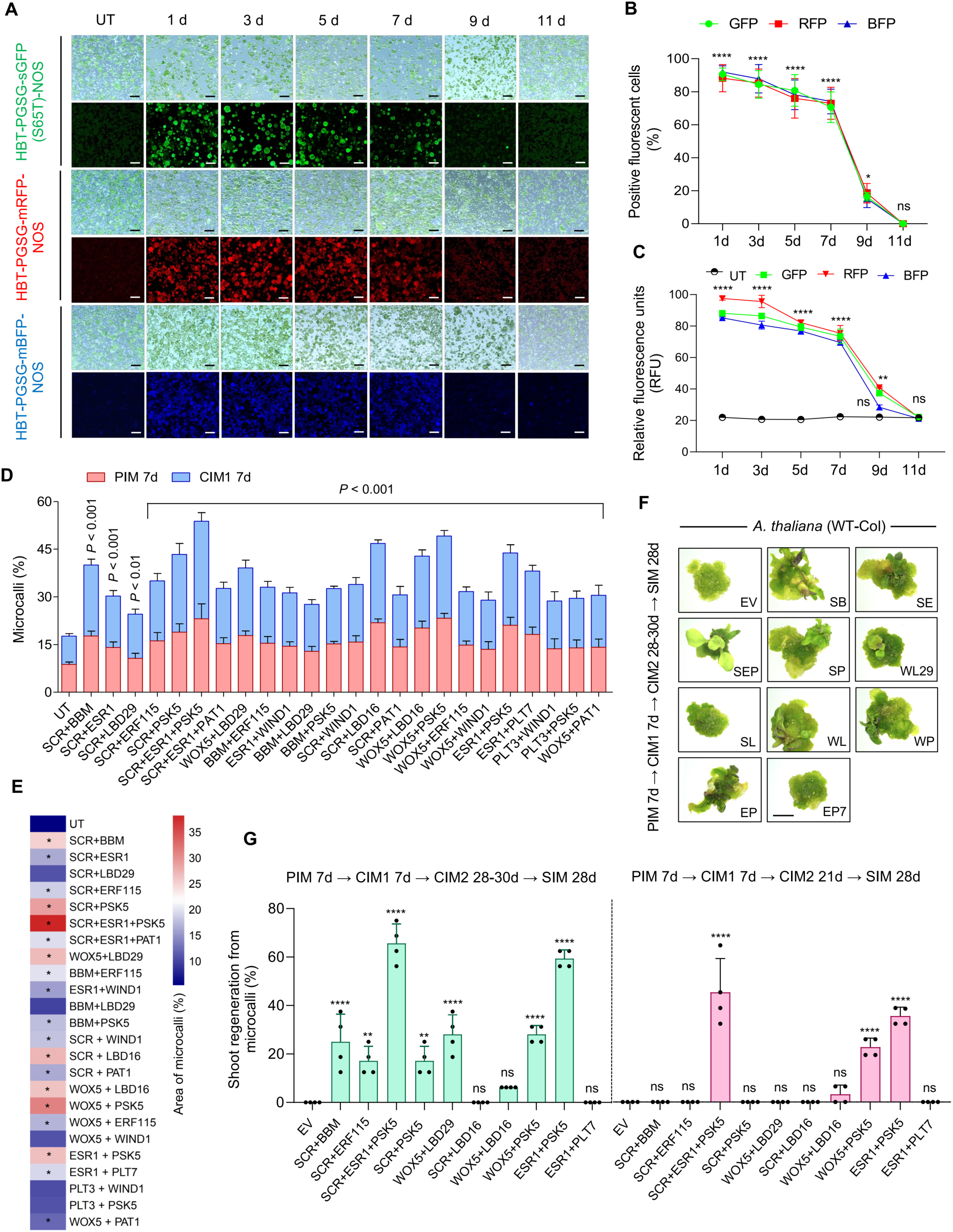
Initial screening of EppTec factors based on the transient expression system. **A**) Bright-field (upper) and fluorescence (lower) micrographs of Arabidopsis mesophyll protoplasts transfected with HBT-PGSG-NOS vectors expressing sGFP, mRFP, or mBFP. Images were captured at the indicated days (d) post-transfection. UT, untransfected control. Scale bars, 50 µm. **B**) Quantification of transfection efficiency over time based on ImageJ analysis of fluorescent protoplasts. **C**) Relative fluorescence units (RFU) measured across time points; the black line indicates mean autofluorescence of the UT control. Data in **(B)** and **(C)** represent mean ± s.d. from three biological replicates (n = 5 each). Statistical comparisons were made to the UT control using two-way ANOVA followed by Dunnett’s multiple comparisons test (\*\*\*\**P* < 0.0001, \*\**P* = 0.0015, \**P* = 0.0109, ns: non-significant). **D**) Percentage of microcalli formation in liquid culture at PIM 7d and CIM1 7d following transfection with the indicated CFs. Microcalli percentage was calculated based on ImageJ analysis of calcofluor white (CW)-stained samples. Data represent mean ± s.d. from five biological replicates (n = 5 each). Statistical significance was determined by one-way ANOVA followed by Dunnett’s multiple comparisons test, comparing each treatment to the UT control at CIM1 7d, and *P* values are indicated above the bars. **E**) Heat map representing the percentage area of microcalli at CIM1 7d, quantified using ImageJ from CW-stained samples. Statistical significance was assessed using one-way ANOVA followed by Dunnett’s multiple comparisons to the UT control. Data represent five biological replicates (n = 5 each). Color scale bar indicates the percentage area of microcalli, and the CFs with *P* < 0.001 are marked with asterisks. **F**) Representative images of regenerated shoots from microcalli under shortened alginate culture conditions. Scale bars, 2 mm. **G**) Percentage of microcalli forming shoots at SIM 28d under two shortened alginate culture timelines. Data represent mean ± s.d. from four biological replicates (n = 16 each). Statistical comparisons were made to the EV control using one-way ANOVA followed by Dunnett’s multiple comparisons test (\*\*\*\**P* < 0.0001, \*\**P* = 0.0014, ns: non-significant). The corresponding culture timelines are indicated in the figure. EV, empty vector control (co-transfection of HBT-PGSG-sGFP(S65T)-NOS, HBT-PGSG-mRFP-NOS, and HBT-PGSG-mBFP-NOS); SB, SCR + BBM; SE, SCR + ERF115; SEP, SCR + ESR1 + PSK5; SP, SCR + PSK5; WL29, WOX5 + LBD29; SL, SCR + LBD16; WL, WOX5 + LBD16; WP, WOX5 + PSK5; EP, ESR1 + PSK5; EP7, ESR1 + PLT7.

### Combinatorial factors promote microcalli development in liquid culture

To investigate whether defined CFs can enhance protoplast regeneration, we selected and cloned Arabidopsis genes shown to be implicated in cellular reprogramming, proliferation, or pluripotency into HBT-PGSG-NOS expression vectors (Xu et al. 2021; Ikeuchi et al. 2018; Kim et al. 2018; Lee et al. 2021; Zhai et al. 2023) (Supplementary Table S1). Each construct was tested for transfection efficiency and sustained expression prior to protoplast culturing. Arabidopsis mesophyll protoplasts were co-transfected with two or three CFs and cultured in liquid PIM to assess division and microcalli formation. In the untransfected (UT) control, < 10% of cells formed microcalli at PIM 7d (Fig. 1D). In contrast, all CFs significantly enhanced microcalli formation, with the SEP [*SCARECROW* (*SCR*) + *ENHANCER OF SHOOT REGENERATION1* (*ESR1*) + *PHYTOSULFOKINE5 PRECURSOR* (*PSK5*)] and WP [*WUSCHEL RELATED HOMEOBOX5* (*WOX5*) + *PSK5*] combinations showing the strongest effects at both PIM 7d and CIM1 7d (Fig. 1D). CW staining confirmed more robust microcalli development in all co-transfected samples relative to the UT control (Supplementary Fig. S3A). Notably, SEP and WP yielded the highest microcalli growth at CIM1 7d (Fig. 1E).

From this initial screen, we identified ten CFs that led to ≥ 2-fold increases in both microcalli percentage and area compared to the UT control (Fig. 1, D and E). Microcalli derived from these CFs were transferred to alginate-embedded CIM2 cultures from CIM1 liquid cultures at d7, and their growth was monitored. By CIM2 at d20∼25, CF-derived microcalli displayed continued growth and diverse morphologies, whereas UT control showed poor or arrested development (Supplementary Fig. S3, B and C). These results establish that transient expression of specific CFs can significantly enhance microcalli formation and growth from Arabidopsis mesophyll protoplasts.

### Three-dimensional alginate scaffolding enables shoot regeneration from microcalli

However, protoplast-derived microcalli transferred from CIM2 30d liquid culture to solid SIM failed to initiate shoot regeneration at SIM 28d. To address this limitation, we introduced a sodium-alginate embedding step for protoplast culture. Protoplasts co-transfected with selected CFs formed earlier and larger microcalli than the empty vector (EV) control at CIM1 7d. These microcalli continued to grow robustly during CIM2 cultures, surpassing EV control in all cases except SL [*SCR* + *LATERAL ORGAN BOUNDARIES-DOMAIN16* (*LBD16*)] and EP7 [*ESR1* + *PLETHORA7* (*PLT7*)] (Supplementary Fig. S4, A and B). Following 28∼30d in CIM2 alginate culture, microcalli were transferred onto SIM. Most CFs promoted efficient shoot regeneration at SIM 28d, with the SEP (SCR + ESR1 + PSK5), EP (ESR1 + PSK5), WP (WOX5 + PSK5), and WL29 (WOX5 + LBD29) groups showing the highest regeneration percentages. SEP yielded the most pronounced effect, reaching ∼65% regeneration, while EV control failed to produce shoots under the same conditions (Fig. 1, F and G, Supplementary Fig. S4, C and D). To further assess regeneration efficiency, we shortened the CIM2 culture period to 20-21d. Under this condition, only SEP, WP and EP transfections supported substantial shoot regeneration, with SEP again performing best (∼45%) (Fig. 1G, Supplementary Fig. S5, A to D). Shoots regenerated from SEP, EP, WP, and WL29 subsequently developed roots on RIM and matured into whole plants (Supplementary Fig. S6, A to C). Further reducing culture duration (PIM 7d ➜ CIM1 7d ➜ CIM2 10d or 14d) abolished shoot regeneration, indicating insufficient microcalli development (Supplementary Figs. S4E and S5E). Together, these results demonstrate that a three-dimensional alginate matrix is essential for reprogrammed microcalli to acquire pluripotency and support robust shoot regeneration, unlike liquid culture alone.

### SEPW and related factor combinations (SEP, EPW, and PWL) support robust transgene-free whole plant regeneration

Given the absence of shoot regeneration in the EV control under the two shortened culture conditions above, we re-assessed shoot regeneration for the UT and EV controls under the standard alginate culture condition (Supplementary Fig. S1D). Under this condition, well-developed microcalli (∼2 mm) with green coloration were obtained from both UT control and EV co-transfected protoplasts at CIM2 30d (Supplementary Fig. S7A), and these microcalli regenerated into shoots, albeit with low efficiencies of ∼17% at SIM 28d (Supplementary Fig. S7, B to D). Toluidine blue staining revealed progressive stages of shoot organogenesis, from meristematic activity observed at SIM 7d to organized shoot primordia at SIM 10d and mature green shoot initiation at SIM 14d (Supplementary Fig. S7E).

To optimize regeneration, we redesigned factor combinations based on initial screening results, assembling three three-factor groups (SEP, EPW, PWL) and one four-factor group (SEPW). Under standard alginate culture, all four combinations promoted stronger microcalli formation, with SEPW showing the highest frequency (∼55%), a ∼4-fold increase over UT and EV controls (Fig. 2A, Supplementary Fig. S8, A to D). These microcalli also yielded superior shoot regeneration, with SEPW achieving a frequency of up to ∼90%, more than five times higher than the UT and EV controls (Fig. 2B, Supplementary Fig. S8, E to H). Regenerated shoots consistently rooted on RIM (Supplementary Fig. S8, I and J).

**Figure 2.**
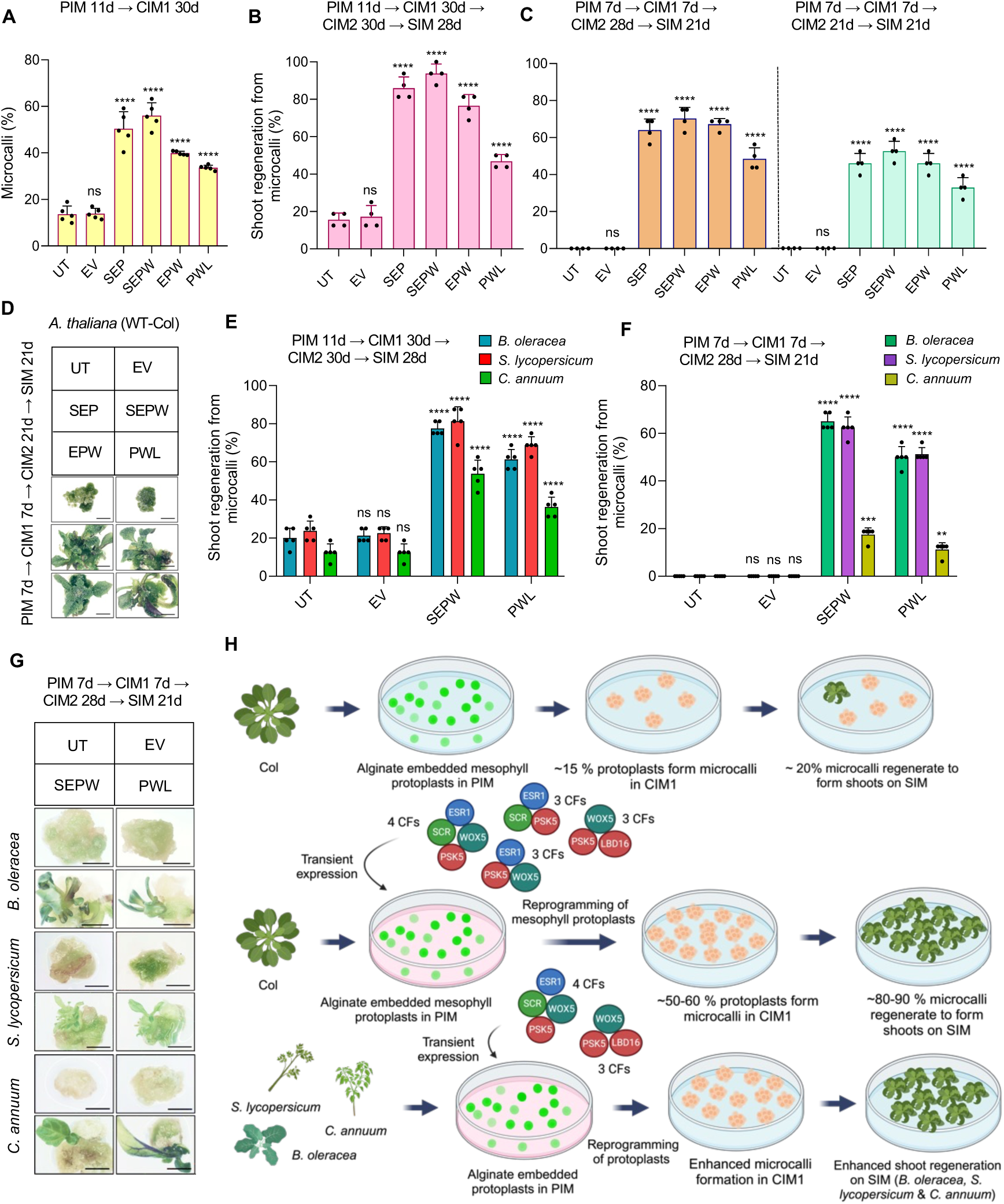
Optimized Arabidopsis CFs promote efficient shoot regeneration from the protoplasts of Arabidopsis and three crop species under standard and shortened alginate culture conditions. **A**) Quantification of microcalli percentage at CIM1 30d following PIM 11d culture, based on ImageJ analysis of CW-stained samples. Data represent mean ± s.d. from five biological replicates (n = 5 each). Statistical comparisons were made to the UT control using one-way ANOVA followed by Dunnett’s multiple comparisons test (\*\*\*\**P* < 0.0001, ns: non-significant). **B)** and **C**) Percentage of microcalli forming shoots in Arabidopsis under standard alginate culture **(B)** and two shortened culture **(C)** conditions. Data represent mean ± s.d. from four biological replicates (n = 16 each). Statistical comparisons were made to the UT control using one-way ANOVA followed by Dunnett’s multiple comparisons test (\*\*\*\**P* < 0.0001, ns: non-significant). **D**) Representative images of the regenerated shoots at SIM 21d from Arabidopsis protoplasts under one shortened culture condition. Scale bars, 2 mm. **E)** and **F**) Percentage of microcalli forming shoots in three crop species under standard **(E)** and one shortened **(F)** alginate culture conditions. Data represent mean ± s.d. from five biological replicates (n = 16 each). Statistical comparisons were made to the UT control using one-way ANOVA followed by Dunnett’s multiple comparisons test (\*\*\*\**P* < 0.0001, \*\*\**P* < 0.001, \*\**P* < 0.01, ns: non-significant). **G**) Representative images of the regenerated shoots from microcalli of three crop species at SIM 21d under one shortened culture condition. Scale bars, 2 mm. **H**) Schematics illustrating a boost in the production of transgene-free regenerated Arabidopsis and crop plants from protoplasts by EppTec using Arabidopsis CFs (EppTec^CF(At)^). The Schematics were created with BioRender.com. The corresponding culture timelines are indicated in each figure panels.

We next evaluated these combinations under two shortened alginate protocols (Supplementary Fig. S1, E and F). All four combinations (SEPW, SEP, EPW, and PWL) consistently outperformed UT and EV controls, which failed to regenerate shoots under these conditions. Among the four CFs, SEPW remained the most effective across both microcalli formation and shoot regeneration (Fig. 2, C and D, Supplementary Figs. S9, A to I, and S10, A to H). All regenerated shoots developed roots (Supplementary Fig. S10, I and J). However, under a further shortened culture condition (PIM 7d ➜ CIM1 7d ➜ CIM2 14d), no shoot regeneration was observed, even with SEP or SEPW, suggesting insufficient pluripotency establishment (Supplementary Fig. S11, A and B). Whole plants were successfully regenerated and acclimated in the soil in all cases (Supplementary Fig. S12, A to F). Regenerated plants from all conditions survived acclimatization and produced viable seeds. Genotyping confirmed the absence of transiently transfected genes in the regenerated plants (Supplementary Fig. S13, A to E), demonstrating that SEPW and related combinations enable efficient, transgene-free regeneration of whole plants from Arabidopsis mesophyll protoplasts.

### EppTec leverages Arabidopsis reprogramming modules to restore regeneration across multiple crop species

To assess whether Arabidopsis CFs can enhance regeneration across species, we tested protoplast-derived regeneration in both closely and distantly related crop species, including *Brassica oleracea* L. cv. Okina, *Solanum lycopersicum* L. cv. MicroTom, and *Capsicum annuum* L. cv. C15 under the standard Arabidopsis protoplast culture protocol (Supplementary Fig. S1D) without any species-specific cultural modifications. Two of the Arabidopsis CFs, SEPW and PWL, were utilized for the co-transfection of protoplasts derived from these crop species. SEPW and PWL co-transfected mesophyll protoplasts of *B. oleracea* and *S. lycopersicum* exhibited superior microcalli development during PIM and CIM1 cultures compared to UT and EV controls (Supplementary Fig. S14, A to D). Although relatively poor microcalli-formation rates were observed at PIM 7d in the mesophyll and hypocotyl protoplasts of *C. annuum* co-transfected with SEPW or PWL (Supplementary Fig. S14E), evidently, there was a higher percentage and improved microcalli development at CIM1 7d and CIM1 30d compared to UT and EV controls, particularly in the hypocotyl protoplasts (Supplementary Fig. S14, F to H). Importantly, SEPW and, to a lesser extent, PWL co-transfections significantly enhanced shoot regeneration from microcalli in all three crop species compared to UT and EV controls (Fig. 2E). Notably, even in *C. annuum*, which exhibited lower initial microcalli formation, both combinations led to marked improvements in shoot regeneration on SIM (Fig. 2E, Supplementary Fig. S15, A and B). Across all three species, regenerated shoots developed roots and whole plants that acclimated successfully in soil (Supplementary Fig. S15, C and D). We next tested SEPW and PWL under the two shortened culture protocols (Supplementary Fig. S1, E and F). Under these conditions, no shoot regeneration occurred in UT or EV controls, while SEPW and PWL induced robust microcalli development and efficient shoot regeneration in all three species (Fig. 2, F and G, Supplementary Figs. S16, A and B, and S17, A to D). All regenerated shoots produced roots and developed into whole plants (Supplementary Figs. S16C and S17E). Genotyping confirmed that the Arabidopsis CFs used for transient transfections were absent from the regenerated plants (Supplementary Figures S18, A to J). These results demonstrate that EppTec using CFs from *Arabidopsis thaliana* (EppTec^CF(At)^) enables transgene-free regeneration in diverse crop species, even under conditions where regeneration is otherwise unsuccessful (Fig. 2H).

### SEPW orthologs enable cross-species protoplast reprogramming and regeneration in rice and a recalcitrant soybean variety

To determine whether SEPW-mediated protoplast reprogramming and regeneration is a conserved process across diverse plant species, we first validated the sustained expression of Arabidopsis and rice factors in rice protoplasts. Transfection with HBT-PGSG-NOS vectors encoding sGFP, mRFP, and mBFP alone or fluorescent-tagged Arabidopsis and rice factors (Supplementary Table S2) showed persistent and robust expression in rice leaf-sheath protoplasts up to 8d post-transfection, similar to the results in Arabidopsis mesophyll protoplasts, indicating the compatibility of the expression system across species (Supplementary Fig. S19, A to E).

This result enabled us to investigate the regeneration potential of both Arabidopsis (AtSEPW and AtPWL) and rice (OsSEPW and OsPWL; Supplementary Table S1) CFs in rice (*Oryza sativa* var. *japonica* cv. Dongjin). Under a standard alginate culture (Supplementary Fig. S19F), co-transfection of the Arabidopsis CFs or rice CFs into rice protoplasts resulted in enhanced microcalli formation compared to UT and EV controls. Notably, OsSEPW and OsPWL combinations induced significantly larger and more proliferative microcalli than AtSEPW and AtPWL combinations, respectively (Fig. 3A, Supplementary Fig. S20, A to D). Histological analysis of microcalli at SIM 6d revealed early signs of cellular differentiation and chloroplast-rich tissues in the CF-transfected microcalli, especially those transfected with rice CFs, which were not observed in UT and EV controls, indicating that rice orthologs initiate cellular differentiation more efficiently than Arabidopsis factors in rice (Fig. 3B). By SIM 10d, shoot-like structures began emerging specifically from microcalli derived from OsSEPW and OsPWL co-transfected protoplasts (Supplementary Fig. S21A), marking the onset of visible shoot organogenesis. In contrast, AtSEPW and AtPWL co-transfections resulted in delayed and intermediate responses, whereas UT and EV controls showed no shoot emergence at SIM 10d. These findings were confirmed at SIM 28d, where OsSEPW co-transfection consistently yielded the highest shoot regeneration efficiency (∼45%), followed by OsPWL, AtSEPW, and AtPWL, with minimal regeneration in UT and EV controls (∼5%) (Fig. 3C). Shoots successfully regenerated roots on RIM (Supplementary Fig. S21B). These results demonstrate that while Arabidopsis CFs can induce reprogramming and regeneration in rice, rice orthologs exhibit better performance. Under a shortened alginate culture protocol (Supplementary Fig. S19G), only OsSEPW and OsPWL co-transfected protoplasts retained the capacity to support microcalli proliferation, cellular differentiation, and whole rice plant regeneration (Fig. 3, D and E, Supplementary Figs. S22, A to C, and S23, A and B). Although AtSEPW and AtPWL showed some effects on promoting microcalli growth and development, they failed to enhance shoot formation, performing comparably to UT and EV controls.

**Figure 3.**
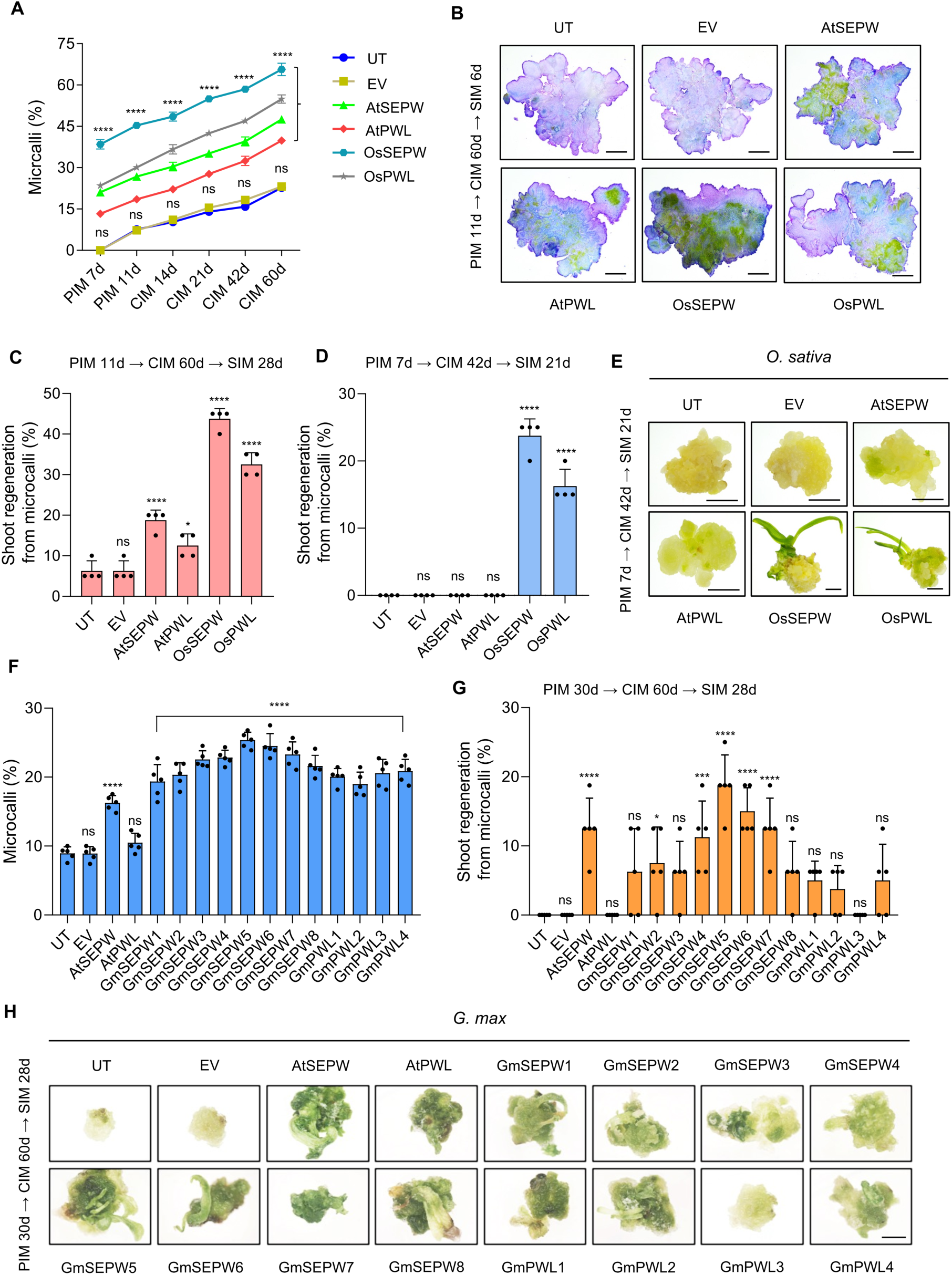
Rice and soybean SEPW/PWL orthologs drive efficient microcalli formation and shoot regeneration under standard and shortened alginate culture conditions. **A**) Quantification of microcalli formation from rice leaf-sheath protoplasts at indicated time points in PIM and CIM cultures, quantified by ImageJ analysis of CW-stained samples. Data represent mean ± s.d. from four biological replicates (n = 5 each). Statistical significance was determined using two-way ANOVA followed by Dunnett’s multiple comparisons test compared to the UT control (\*\*\*\**P* < 0.0001, ns: non-significant). **B**) Bright-field images of toluidine blue-stained sections from developing rice microcalli at SIM 6d, showing early tissue organization. Scale bars, 500 µm. **C)** and **D**) Percentage of rice microcalli forming shoots at SIM 28d under standard alginate culture **(C)** and at SIM 21d under a shortened alginate culture condition **(D)**. Data represent mean ± s.d. from four biological replicates (n = 20 each). Statistical significance was assessed using one-way ANOVA followed by Dunnett’s multiple comparisons test compared to the UT control (\*\*\*\**P* < 0.0001, \**P* < 0.05, ns: non-significant). **E**) Representative images of regenerated rice shoots at SIM 21d under the shortened alginate culture. Scale bars, 2 mm. **F**) Percentage of microcalli formed from soybean protoplasts at PIM 30d, quantified from CW-stained samples. Data represent mean ± s.d. from five biological replicates (n = 5 each). Statistical comparisons were made to the UT control using one-way ANOVA followed by Dunnett’s multiple comparisons test (\*\*\*\**P* < 0.0001, ns: non-significant). **G**) Percentage of soybean microcalli forming shoots at SIM 28d under standard alginate culture. Data represent mean ± s.d. from five biological replicates (n = 16 each). Statistical significance was assessed using one-way ANOVA followed by Dunnett’s multiple comparisons test compared to the UT control (\*\*\*\**P* < 0.0001, \*\*\**P* = 0.0004, \**P* = 0.0403, ns: non-significant). **H**) Representative images of regenerated soybean shoots at SIM 28d under standard alginate culture. Scale bar, 2 mm. The corresponding culture timelines are indicated in the figure.

Applying the same approach in a recalcitrant soybean variety (*Glycine max* var. Williams 82) revealed that the GmSEPW5 (Supplementary Tables S1 and S2) was the best combination to robustly induce both microcalli development and shoot regeneration under a standard alginate culture, where no shoot regeneration occurred in UT or EV controls (Fig. 3, F to H, Supplementary Fig. S24, A and B). AtSEPW co-transfection showed a moderate effect on microcalli formation and shoot regeneration in soybean mesophyll protoplasts. Shoots successfully regenerated roots on RIM (Supplementary Fig. S24C). When soybean protoplasts were subjected to a shortened culture protocol (Supplementary Fig. S25A), only GmSEPW5 and GmSEPW6 co-transfected protoplasts supported shoot regeneration, although AtSEPW and all soybean CFs showed effectiveness until the stage of microcalli development (Supplementary Fig. S25, B to E). In summary, the results above demonstrate that SEPW/PWL-mediated reprogramming is evolutionarily conserved across dicot and monocot, and the use of species-specific or evolutionarily less diversified SEPW/PWL orthologs clearly further enhances the efficacy of EppTec, offering a powerful strategy to drive regeneration in diverse and challenging plant species.

### SEPW triggers epigenomic reprogramming and cell-fate transition

To investigate the molecular basis of SEPW-induced protoplast reprogramming, we first examined global histone modifications at 3d post-transfection. Immunostaining of Arabidopsis and rice nuclei transfected with AtSEPW and OsSEPW, respectively, revealed a significant increase in histone H3 lysine 4 tri-methylation (H3K4me3), a representative activation mark, and a decrease in H3K27me3, a representative repression mark (Fig. 4, A and B), indicating an epigenetic shift toward a transcriptionally permissive chromatin state. To assess the downstream consequence of this epigenetic shift, we performed single-cell RNA-sequencing (scRNA-seq) on AtSEPW-transfected Arabidopsis mesophyll protoplasts (Supplementary Fig. S26, A to C). UMAP projection identified an SEPW-specific reprogrammed cell cluster, which was absent in mock control (Fig. 4C), indicating an SEPW-induced cell-fate transition. Cell cycle analysis showed that these reprogrammed cells were predominantly in the S phase, in contrast to mesophyll-cell clusters, which were enriched in G0-like/quiescent cells (Fig. 4D), suggesting cell cycle re-entry during reprogramming. Heatmap analysis of top upregulated genes in the reprogrammed cell cluster highlighted cell cycle and transcriptional signatures associated with SEPW-induced reprogramming (Fig. 4E). Not surprisingly, the transcripts of the four genes used for transfection (*SCR*, *ESR1*, *PSK5*, and *WOX5*) were also highly enriched in this cell cluster (Supplementary Fig. S26D). Gene ontology analysis of SEPW-induced genes revealed enrichment in biological processes related to translation, cell cycle reactivation, response to wounding, auxin response, and others (Supplementary Fig. S26E). Consistent with the scRNA-seq evidence of cell cycle activation, 5-ethynyl 2’-deoxyuridine (EdU) incorporation assays showed a marked enrichment of S-phase nuclei in SEPW-transfected protoplasts compared with controls, confirming that SEPW promotes DNA synthesis and cell cycle re-entry (Fig. 4, F and G). Taken together, these findings demonstrate that SEPW factors induce epigenomic reprogramming and establish a transcriptional state associated with cellular reprogramming.

**Figure 4.**
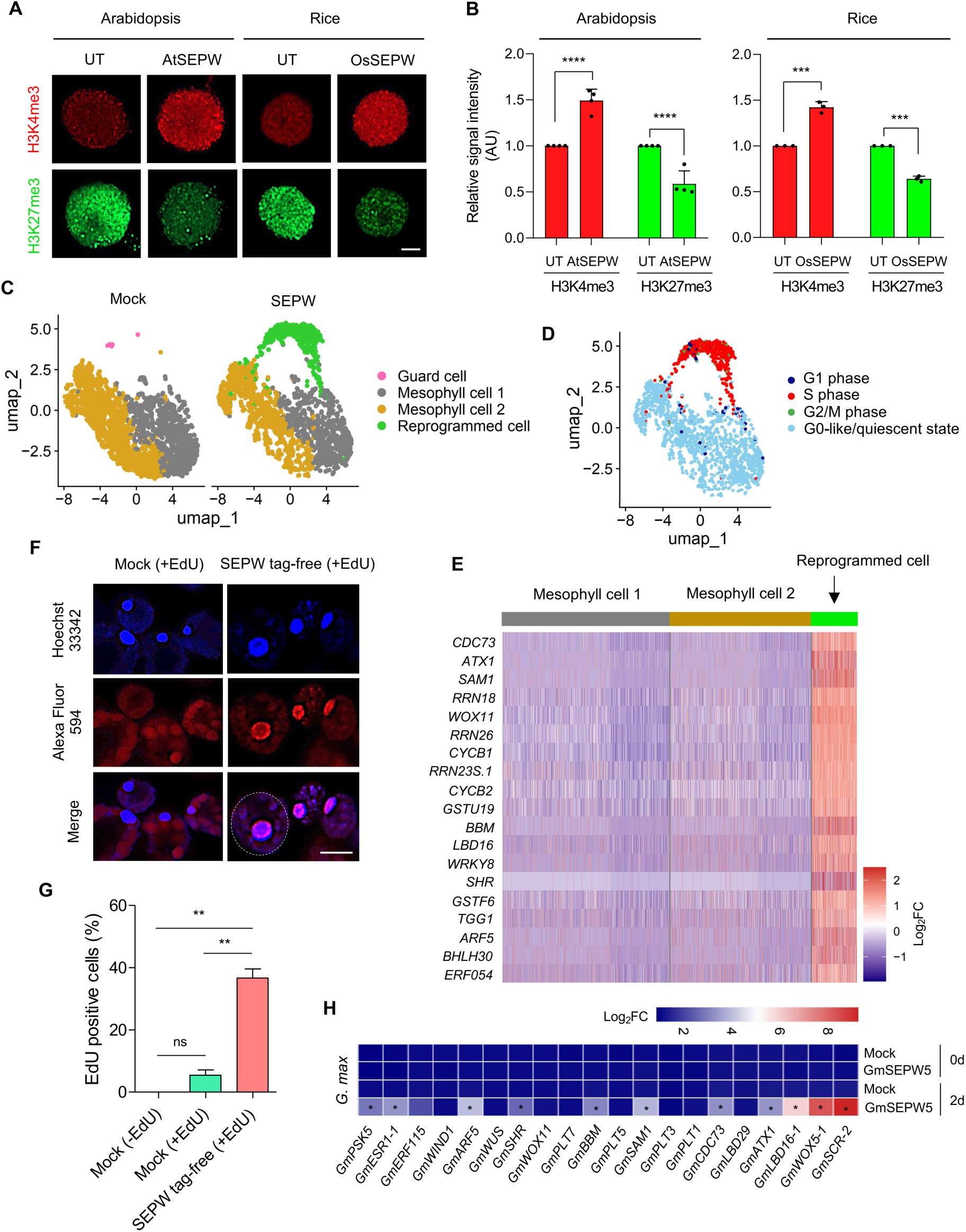
SEPW induces epigenomic reprogramming and cell-fate transition. **A**) Immunostaining of nuclei isolated from Arabidopsis and rice protoplasts at 3d post-transfection, showing changes in histone methylation states upon SEPW expression. Representative confocal images show nuclear signals for H3K4me3 (red) and H3K27me3 (green) in UT and SEPW-transfected protoplasts. Scale bar, 2 µm. **B**) Quantification of H3K4me3 and H3K27me3 signal intensities in Arabidopsis and rice protoplast nuclei, normalized using ZEN software. Data represent mean ± s.d. from four (Arabidopsis) and three (rice) independent biological replicates (n = 10 each). AU is the arbitrary unit. Statistical significance was assessed using two-tailed unpaired *t*-tests compared to the UT control (\*\*\*\**P* < 0.0001, \*\*\**P* < 0.001). **C**) UMAP visualization of scRNA-seq profiles from mock and SEPW-transfected Arabidopsis mesophyll protoplasts. Cell clusters are annotated as Guard cell, Mesophyll cell 1, Mesophyll cell 2, and Reprogrammed cell. **D**) Cell cycle phase classification of SEPW-transfected protoplasts. A majority of reprogrammed cells were assigned to the S phase, while mesophyll cell clusters showed an enrichment of G0-like/quiescent cells. **E**) Heatmap showing scaled expression of selected cell cycle-, pluripotency-, and reprogramming-associated genes across Mesophyll cell 1, Mesophyll cell 2, and Reprogrammed cell clusters. Expression values are z-score normalized. **F**) Confocal imaging of EdU labeling in Arabidopsis mesophyll protoplasts at 2d after SEPW (tag-free) transfection. Nuclei were counterstained with Hoechst 33342 (blue), and EdU incorporation was visualized with Alexa Fluor 594 (red). Representative fields show a strong nuclear EdU signal in SEPW (tag-free) transfected protoplasts compared with the mock positive control. Scale bar, 10 µm. The white dotted circle in the merge panel highlights a single protoplast. **G**) Quantification of the percentage of EdU⁺ nuclei in SEPW (tag-free) transfected protoplasts along with the mock controls. Data represent mean ± s.d. from two independent biological replicates (n = 5 each). Statistical significance was assessed using one-way ANOVA followed by Turkey’s multiple comparisons test compared with mock controls (\*\**P* = 0.0011, ns: non-significant). **H**) Heat map showing relative expression levels of endogenous reprogramming-, pluripotency-, and cell cycle-associated genes in soybean mesophyll protoplasts transfected with GmSEPW5 at the indicated time points. Gene expression was normalized to *GmActin4*. Data represent means of three independent biological replicates (n = 3 each). Statistical significance for each gene comparison was assessed using two-tailed unpaired *t*-tests compared to the mock control; genes with *P* < 0.001 are marked with asterisks. Color scale indicates log₂-transformed expression values.

To determine whether the SEPW-induced transcriptional program is conserved beyond Arabidopsis, we analyzed soybean mesophyll protoplasts 2d after GmSEPW5 transfection and evaluated the expression of endogenous soybean genes that are homologous to the Arabidopsis genes upregulated by AtSEPW transfection. RT-qPCR analysis revealed significant upregulation of these endogenous soybean genes in GmSEPW5-transfected cells compared to mock controls (Fig. 4H), implying a conserved transcriptional framework underpinning protoplast reprogramming. In summary, the data above suggest that the SEPW module might engage a shared core regulatory program across diverse species, supporting its broad or universal utility for enabling reprogramming and regeneration.

### Transient expression of SEPW induces stable and long-term reprogramming into a pluripotent state

To investigate the molecular basis of SEPW-induced regeneration, we used homozygous *pWOX5::nYFP* and *pSCR::SCR-mGFP* stable transgenic Arabidopsis plants (Supplementary Fig. S27, A and B), which mark endogenous expression of pluripotency regulators *WOX5* and *SCR*, respectively (Kim et al. 2018). Mesophyll protoplasts isolated from these plants were co-transfected with tag-free SEPW, and expression of the pluripotency markers (i.e., nYFP and SCR-mGFP reporters) was monitored over time. Robust nuclear expression of both nYFP and SCR-mGFP was detected from d1 and persisted for d9 post-transfection in SEPW (tag-free)-transfected protoplasts, while no signal was observed in the UT control (Fig. 5, A to D). In less than 3% of UT protoplasts, low-level background expression was observed at 6 hours (hr), likely due to mechanical stress from protoplast isolation (Xu et al. 2021). At d11, expression declined in non-dividing cells, particularly in protoplasts that failed to form microcalli. In contrast, microcalli derived from SEPW (tag-free)-transfected protoplasts maintained strong expression of both pluripotency markers throughout PIM, CIM1, and CIM2 phases, and even into early SIM culture (SIM 3d) (Fig. 5, E to G and Supplementary Fig. S27, C and D). Nearly 80% of SEPW-derived microcalli consistently expressed both nYFP and SCR-mGFP during this period, while no sustained expression was observed in UT-derived microcalli. These surprising results indicate that transient expression of SEPW initiates a stable reprogramming cascade in mesophyll protoplasts, leading to long-term activation of key pluripotency genes. This transcriptional memory likely underlies the enhanced regenerative capacity observed in SEPW-transfected protoplasts and may represent a mechanistic basis for the establishment of pluripotent cell and microcallus identity.

**Figure 5.**
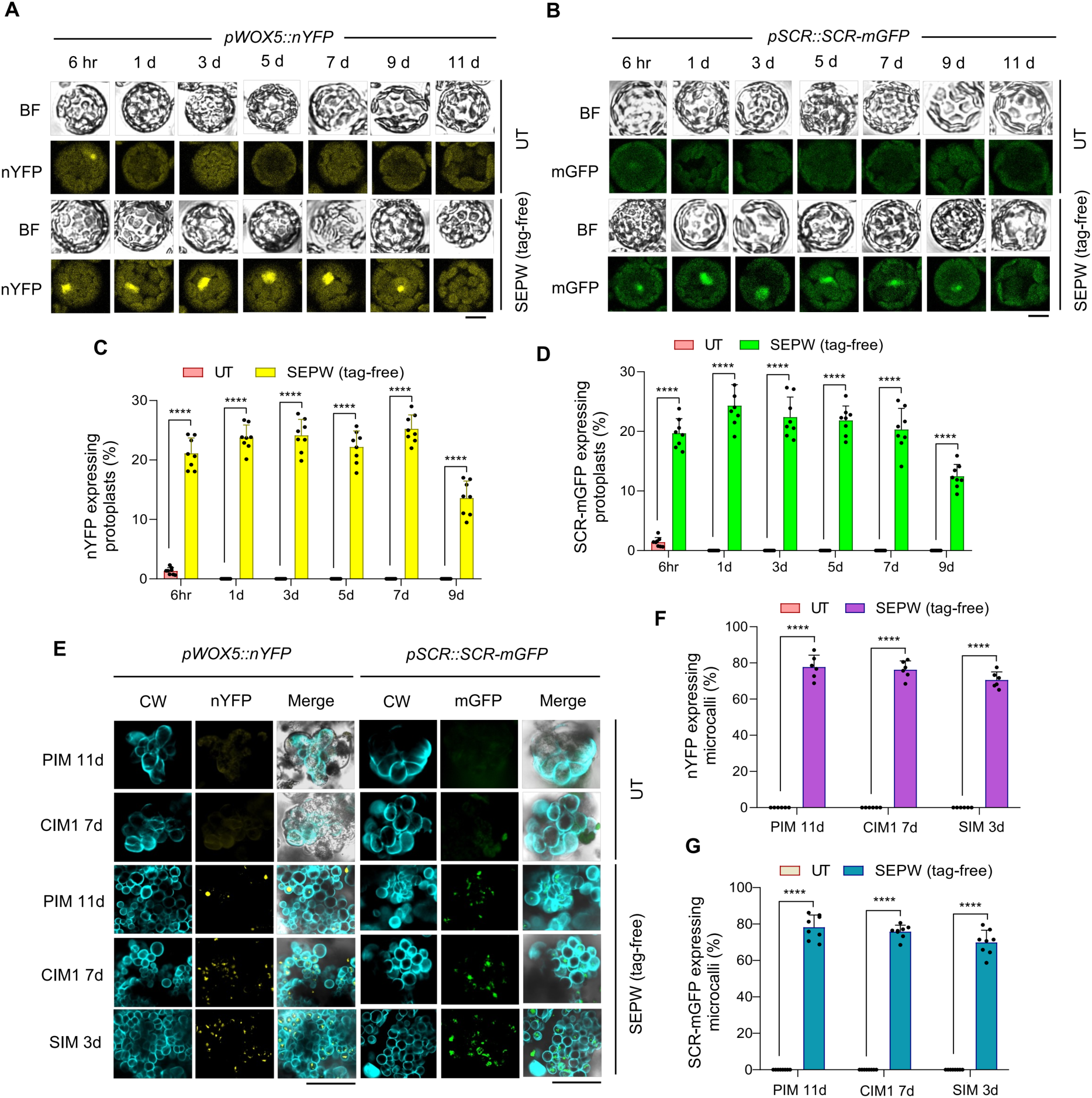
Transient transfection of SEPW induces long-term induction of pluripotency marker genes. **A)** and **B**) Confocal micrographs showing nuclear expression of nYFP **(A)** and SCR-mGFP **(B)** in non-dividing Arabidopsis mesophyll protoplasts following SEPW co-transfection. Protoplasts were isolated from transgenic plants homozygous for *pWOX5::nYFP* **(A)** or *pSCR::SCR-mGFP* **(B)**, and transfected with fluorescence tag-free SEPW constructs. Images were acquired at indicated time points (6hr to 11d) post-transfection under bright-field (BF) and fluorescence channels. Scale bars, 10 µm. **C)** and **D**) Percentage of non-dividing protoplasts expressing nYFP **(C)** or SCR-mGFP **(D)**, quantified at indicated time points. Data represent mean ± s.d. from eight biological replicates (n = 4 each). Statistical comparisons were made to the UT control using two-way ANOVA followed by Dunnett’s multiple comparisons test (\*\*\*\**P* < 0.0001). **E**) Confocal micrographs of developing and dividing microcalli showing nuclear expression of nYFP and SCR-mGFP following SEPW co-transfection. Protoplasts isolated from *pWOX5::nYFP* or *pSCR::SCR-mGFP* plants were embedded in alginate and cultured sequentially in PIM for 11d, CIM1 for 30d, CIM2 for 30d, and SIM for 3d. CW-stained microcalli were imaged at the indicated stages. Scale bars, 300 µm. **F)** and **G**) Percentage of microcalli expressing nYFP **(F)** or SCR-mGFP **(G)** at indicated time points. Data represent mean ± s.d. from six biological replicates (n = 5 each) for **(F)** and eight biological replicates (n = 4 each) for **(G)**. Statistical comparisons were made to the UT control using two-way ANOVA followed by Dunnett’s multiple comparisons test (\*\*\*\**P* < 0.0001).

## Discussion

Plants and animals can regenerate with varying degrees of undifferentiated or partially differentiated stem cells that are primed to acquire pluripotency in response to internal or external stimuli. Currently, there are limited reports on the regeneration of Arabidopsis protoplasts, indicating constrained shoot regeneration efficiency (Dovzhenko et al. 2003; Jeong et al. 2021; Damm and Willmitzer 1988; Siemens et al. 1993). An efficient vector system and viable protoplasts with high transfection efficiency are crucial for the successful use of the protoplast transient-expression system (Liu et al. 2022). In this study, we introduce a novel approach, EppTec, that demonstrates a remarkable enhancement in protoplast regeneration in Arabidopsis and diverse crop species by transiently expressing Arabidopsis CFs and their orthologs. Advancements in liquid culture techniques have increased the rate of protoplast division and enabled temporal transcript profiling during the early events leading to totipotency (Chupeau et al. 2013). Although the co-transfection of Arabidopsis CFs facilitated early protoplast division and promoted increased microcalli formation in a liquid medium, microcalli did not seem to develop proper pluripotency, as revealed by the lack of shoot regeneration ability. Effective tissue regeneration requires intricate coordination between cellular reprogramming and cell cycle progression, which is likely mediated by cell-to-cell communication within structured tissues. Thus, liquid culture systems may not entirely mimic the 3D microenvironment of natural tissues (Luo et al. 2015; Ikeuchi et al. 2016). When EppTec was combined with a protoplast culture in sodium alginate (Luo et al. 2015), a 3D microenvironment was created, which greatly facilitated cellular reprogramming and the acquisition of pluripotency. Future studies aimed at understanding how EppTec components reprogram plant cells and enhance protoplast regeneration potential are warranted. Such studies hold promise for advancing our understanding of the reprogramming processes leading to pluripotency acquisition and for revolutionizing agriculture and plant biotechnology by accelerating plant breeding and crop improvement through genetic modification.

Although the microscopic and alginate-embedded nature of the regenerating cells/tissues presents a significant technical challenge for unraveling the intricate mechanistic details at this stage, elucidating the biochemical roles of individual or combined factors used in EppTec is of substantial importance and value. One notable aspect is the consistent presence of *PSK5* in our four– and three-factor EppTec combinations, which displayed the best performances in enhancing microcalli formation and shoot regeneration. *PSK5* encodes a precursor polypeptide that undergoes processing and modification to form a sulfated pentapeptide (Yang et al. 2000). PSK5 has been established as a potent inducer of cell proliferation in tissue culture, underscoring its crucial role in regenerative processes (Yang et al. 2000; Kielkowska et al. 2019). One possible scenario is that the accelerated cell division driven by PSK5 might enhance stochastic chromatin accessibility, which in turn promotes the targeting of transcriptional regulators, SCR, ESR1, WOX5, and LBD16, used in combination with PSK5 for EppTec. These transcriptional regulators may act as pioneer factors, inducing genes required for cell-lineage specification, leading to pluripotent stem cells and stem-cell niches within developing microcalli. *SCR* and *WOX5* seem to be downstream of SEPW. These two genes are known to play pivotal roles in the establishment and maintenance of the root stem cell niche during embryonic root development and the acquisition of pluripotency in the callus (Kim et al. 2018; Shimotohno et al. 2018). Therefore, a likely scenario is that SCR and WOX5 may self-regulate in a positive feed-forward way and also serve as key pioneer factors. In the context of advancing cellular reprogramming in plants, a crucial avenue for future investigation revolves around elucidating detailed mechanisms by which the factors used for EppTec and other pioneer factors can induce a pluripotent state in protoplasts. Drawing parallels to the paradigm of Yamanaka factors (Oct3/4, Sox2, c-Myc, and Klf4) in generating induced pluripotent stem cells (iPSCs) in animals under embryonic stem-cell culture conditions (Takahashi et al. 2006), a similar comprehensive understanding should be sought in plant biology, and EppTec-based approaches may provide new opportunities in this regard.

Although protoplast regeneration poses a formidable challenge for numerous crop species (Jain et al. 1995; Tan et al. 1987; Nishio et al. 1989; Kiełkowska et al. 2012; Sandgrind et al. 2012), EppTec efficiently enhances protoplast regeneration from recalcitrant species of *C. annuum* and *G. max* (Saxena et al. 1981; Diaz et al. 1988; Donato et al. 1989; Prakash et al. 1997; Wei et al. 1988; Gamborg et al. 1983). Thus, EppTec is believed to boost protoplast regeneration across a broad range of plant species. Notably, orthologous SEPW factors from rice and soybean outperformed their Arabidopsis counterparts within their respective species by accelerating microcalli growth and shoot regeneration, revealing a conserved yet tunable mechanism underlying cellular reprogramming. Mechanistically, SEPW initiates epigenomic reprogramming, marked by elevated H3K4me3 and reduced H3K27me3, resulting in a permissive chromatin environment. Our single-cell transcriptomics revealed a SEPW-induced new cell population enriched for cell cycle progression and reprogramming, while lineage reporters demonstrated sustained activation of pluripotency markers (*WOX5* and *SCR*) during microcalli progression. These findings indicate that SEPW establishes a transcriptionally primed, pluripotent-like state *via* stable reprogramming. Unlike hormone-based approaches, EppTec provides a modular, ortholog-adaptable framework to unlock totipotency across diverse species.

In numerous crop species, the production of transgenic or genome-edited plants relies on the successful regeneration of plants from transformed tissues or edited protoplasts. Despite its necessity for genetic transformation or genome editing, regeneration often acts as a major bottleneck, outweighing the challenges associated with stable DNA integration or editing (Altpeter et al. 2016). EppTec would assist in breaking this bottleneck and enhancing the efficiency and target-species range of plant genome or epigenome engineering. Furthermore, EppTec may also facilitate the regeneration of haploid and hybrid protoplasts. These improvements may streamline the production of genetically modified plants with desired traits, including haploid or hybrid plants, thereby accelerating plant breeding efforts (Reed and Bargmann 2021). In addition, EppTec may also help efforts to preserve endangered plant species by promoting their regeneration in a transgene-free way. The ability to overcome historical challenges in protoplast regeneration not only enhances our understanding of the excellent plasticity or regeneration potential of plants but also provides a robust platform for future innovations in plant biotechnology/agriculture and conservation biology.

## Materials and methods

### Vectors and plasmids used for EppTec

We initially devised highly efficient vector systems for transient gene expression in protoplasts, using the NEBuilder HiFi DNA Assembly Reaction protocol (New England Biolabs, E2621S). The mother vector, HBT-sGFP(S65T)-NOS (Yoo et al. 2007), served as the precursor, which was modified by the addition of a DNA sequence encompassing multiple cloning sites (MCS) containing *Bam*HI and *Xma*I*/Sma*I sites. The HBT vector contains a modified CaMV 35S promoter (35SPPDK) with dual enhancers, including elements from the maize PPDK promoter, which enhances transcriptional activity in plant cells (Yoo et al. 2007). Four amino acids (pro-gly-ser-gly) were encoded downstream of the 35SPPDK promoter, resulting in the creation of the modified HBT-PGSG-sGFP(S65T)-NOS vector. To generate the HBT-PGSG-mRFP-NOS or HBT-PGSG-mBFP-NOS vectors, the sGFP(S65T)-encoding sequence in the HBT-PGSG-sGFP(S65T)-NOS vector was substituted with the mRFP– or mBFP-encoding sequence, respectively. The HBT-PGSG-sGFP(S65T)-NOS vector was initially linearized using *Stu*I digestion and purified using a DNA purification kit (Intron Biotechnology, 17290). High-fidelity *pfu*-X DNA polymerase (SolGent, SPX16-R250) was employed to amplify both the vector and the *sGFP*/*mRFP*/*mBFP* inserts, utilizing overlapping primers (Supplementary Table S2). To eliminate the plasmid template, the PCR product was digested with *DpnI* and purified using a PCR purification column (Intron Biotechnology, 17290). The NEBuilder HiFi DNA assembly reaction was initiated after removing the plasmid template. Ligation involved the PCR-amplified vector backbone and the *sGFP*/*mRFP*/*mBFP* inserts, as per the manufacturer’s protocol. The resulting ligation mixture was transformed into *E. coli* DH5α and colonies resistant to ampicillin were subjected to further screening through colony PCR using specific primers designed for the 35SPPDK promoter and the *sGFP*/*mRFP*/*mBFP* regions (Supplementary Table S2). Positive clones were ultimately confirmed through sequencing (Supplementary Table S2). For gene cloning into the designated HBT-PGSG-sGFP/mRFP/mBFP-NOS vectors, first-strand cDNA was synthesized from total RNA using the RevertAid First Strand cDNA Synthesis Kit (Thermo Fisher Scientific, K1621). Specific coding sequences (CDSs) were then amplified by PCR using overlapping primers (Supplementary Table S2). The subsequent HiFi DNA assembly involved PCR-amplified CDS and *Sma*I-digested vector. Genes such as *SCR*, *ESR1*, *PSK5*, and *WOX5* were also cloned into the vectors without *sGFP*/*mRFP*/*mBFP* tags for tag-free experiments. The desired genes and vector backbones (without tags) were amplified (Supplementary Table S2) and assembled using the NEBuilder HiFi DNA Assembly protocol. The final clones were validated by sequencing (Supplementary Table S2).

### Identification and selection of orthologs in rice and soybean

To identify functional orthologs of Arabidopsis SEPW/PWL factors (SCR, ESR1, PSK5, WOX5, and LBD16) in rice (*Oryza sativa*) and soybean (*Glycine max*), we performed multiple sequence alignment and phylogenetic analyses using Clustal Omega and MEGA11 software. Protein sequences of Arabidopsis factors were used as queries to retrieve putative homologs from the Rice Genome Annotation Project database (https://rice.uga.edu) and the SoyBase repository (https://www.soybase.org/). Structural predictions were generated using AlphaFold2, and candidates showing conserved α-helical folding and domain architecture relative to Arabidopsis proteins were prioritized (Jumper et al. 2021). Gene expression profiles from the rice expression database were used to shortlist orthologs with high transcript abundance in callus tissue. Accordingly, total RNA was isolated from 10-d-old scutellum-derived rice callus, and first-strand cDNA was synthesized as described in the previous section. Soybean cDNA was synthesized from total RNA isolated from 7-d-old hypocotyl-derived callus. CDSs of the selected genes from rice and soybean were cloned into the HBT-PGSG-sGFP/mRFP/mBFP-NOS vectors for transient transfection into the protoplasts (Supplementary Table S2).

### Plasmid preparation by CsCl density gradient ultracentrifugation

Plasmids were purified from *E. coli* DH5α cells cultured in 250 mL LB medium (Condalab, CAT. 1551.00) containing ampicillin using the alkaline lysis method (Birnboim and Doly 1979). The extracted DNA was resuspended in 2 mL TE buffer (10 mM Tris-HCl, pH 8.0, 1 mM EDTA), followed by the addition of 2.2 g CsCl (Thermo Fisher Scientific, BP1595-500) and 160 µL ethidium bromide (10 mg/mL stock; Sigma-Aldrich, E8751). After CsCl had fully dissolved, the solution was transferred into polycarbonate ultracentrifuge tubes (Beckman Coulter, TLA110) and centrifuged at 65,000 rpm for 14∼16 hr at 22°C using an Optima MAX-XP ultracentrifuge (Beckman Coulter). The supercoiled form of plasmid was carefully extracted using a syringe and subsequently washed with 1-butanol (Sigma, B7906-500ML) four to five times until the solution cleared. The purified plasmid was concentrated through ethanol precipitation and quantified using Nanodrop (GE Healthcare).

### Plant materials and growth conditions

*Arabidopsis thaliana* accession Columbia-0 (Col) seeds were surface-sterilized by washing 5∼6 times in 0.1% Triton X-100 (Thermo Fisher Scientific, A16046.AP) dissolved in 70% ethanol (Merck, 1.00983.1011), followed by two washes with 90% ethanol. Sterilized seeds were air-dried on sterile filter paper and sown on solid MS medium (Supplementary Table S3). Seeds were stratified at 4°C in the dark for 2∼3d and then transferred to a growth chamber under long-day conditions (16 hr light/8 hr dark) at 22°C. For *Brassica oleracea* L. cv. Okina, *Solanum lycopersicum* L. cv. MicroTom, and *Capsicum annuum* L. cv. C15, seeds were sterilized similarly, then treated with 3% sodium hypochlorite (Sigma-Aldrich, MSDS-105614) for 2 min, followed by 4∼5 washes with sterile water. Seeds were dried on sterile filter paper and sown on either solid MS medium (*B. oleracea*) or half MS medium supplemented with 1.5% sucrose (*S. lycopersicum* and *C. annuum*). After stratification at 4°C for 4d in the dark, *B. oleracea* was germinated at 22°C, while *S. lycopersicum* and *C. annuum* were germinated at 25°C, all under long-day photoperiod. *Oryza sativa* (var. *japonica* cv. Dongjin) and *Glycine max* (var. Williams 82) seeds were sterilized by repeated washing (5∼6 times) with the same ethanol-Triton X-100 solution, followed by treatment with 3% sodium hypochlorite for 2 min and thorough rinsing (4∼5 times) with sterile water. Rice seeds were sown directly on solid MS medium and incubated under long-day photoperiod at 25°C. Soybean seeds were placed on an autoclaved cotton bed soaked with half-strength liquid MS medium containing 3% sucrose, germinated in the dark for 7d, and subsequently grown under long-day photoperiod at 28°C for an additional 7d.

### Protoplast isolation

Protoplasts were isolated as previously described with minor modifications (Yoo et al. 2007). Approximately 100∼150 leaves from 15-d-old *Arabidopsis thaliana* (Col) plants grown on MS medium were cut and incubated in 10 mL enzyme solution for 5∼6 hr at 22°C in the dark to release mesophyll protoplasts. The digestion mixture was diluted 1:1 with W5 solution and filtered through a 40 μm cell strainer (SPL Life Sciences, 93040) to remove undigested tissue. Protoplasts were pelleted by centrifugation at 600 rpm for 10 min, washed twice with W5 solution, and resuspended in a reduced volume of MMG solution (Supplementary Table S3). Cell density was determined using a hemocytometer under a light microscope (Carl Zeiss, Axioskop 40) and adjusted to ∼2 × 10⁵ cells/mL in MMG solution for transfection or culture. Protoplasts from *Brassica oleracea* (leaf), *Solanum lycopersicum* (cotyledon), and *Capsicum annuum* (leaf or hypocotyl) were isolated from 2∼3-week-old seedlings using the same procedure. For rice (*Oryza sativa*), protoplasts were isolated from 12-d-old leaf base tissue as described above, with additional purification by 21% (w/v) sucrose density gradient centrifugation, followed by washing with W5 solution and resuspension in MMG buffer. For soybean (*Glycine max*), protoplasts were isolated from unifoliate leaves of 15-d-old seedlings following the same Arabidopsis-based protocol. All solutions used for protoplast isolation, transfection, and culture are listed in Supplementary Table S3.

### DNA-PEG-calcium mediated protoplasts transfection

Numerous transfection conditions have been extensively studied, each yielding different success rates depending on the nature of protoplasts (Liu et al. 2022; Stajic and Kunei 2023; Yang et al. 2024). In this study, PEG-mediated transfection was performed in Arabidopsis and crop species following a previously described method with minor modifications (Yoo et al. 2007). Protoplasts were pre-incubated in the dark at 22°C for 1 hr prior to transfection. Approximately 10 µg of supercoiled plasmid DNA (purified using CsCl density gradient ultracentrifugation) was mixed with 100 µL of protoplast suspension in MMG solution at a density of ∼2 × 10⁴ cells/mL. A total of 150 µL of 20% PEG-4000 (Fluka, 81240) solution was added to the mixture, and the suspension was incubated at room temperature in the dark for 10∼15 min. The transfection was terminated by adding an equal volume of W5 solution, followed by centrifugation at 500 rpm for 5 min. Protoplasts were washed once with PIM and resuspended in 500 µL of PIM for imaging and downstream culture (Supplementary Table S3).

### Protoplast or microcalli staining

Protoplast viability was assessed using fluorescein diacetate (FDA; Invitrogen, F1303) staining, as described previously (Jones and Senft 1985). FDA staining solution was prepared by diluting 20 µL of a 5 mg/mL FDA stock into 1 mL of 0.8 M D-mannitol (Ducefa Biochemie, M0803.1000). For staining, 5 µL of FDA solution was added to 10 µL of protoplast suspension, incubated for 2 min at room temperature, and washed with W5 solution (Supplementary Table S3) prior to imaging. Calcofluor white (CW; Sigma-Aldrich, 18909-100ML-F) was used to stain dividing protoplasts and developing microcalli in PIM or CIM cultures (Monthony and Jones 2024). For Arabidopsis liquid cultures, an equal volume of protoplast suspension from PIM or CIM1 was mixed with 1:10 diluted CW for 2 min, followed by a W5 wash prior to imaging. For alginate-embedded cultures, a small block of the alginate layer was excised, stained in CW for 2 min in a separate Petri dish (60 × 15 mm, SPL Life Sciences, 11060), and imaged using a fluorescence stereomicroscope (Carl Zeiss, AxioZoom V16) or confocal microscope (Carl Zeiss, LSM700) after washing with W5. Propidium iodide (PI; Sigma-Aldrich, P4170) was used to visualize dead protoplasts or cell wall integrity. A 1 mg/mL PI stock solution was used to stain tissues for 1 min, followed by washing with sterile water. Imaging was performed using a confocal microscope (Carl Zeiss, LSM700).

### Protoplast culture

Various support matrices were tested for protoplast culture, including low-melting agarose (Sigma-Aldrich, A9414), VitroGel (TheWell Bioscience, VHM01), phytoagar (Ducefa Biochemie, P1003.1000), and sodium alginate (Sigma-Aldrich, A2158). For liquid culture, protoplasts were plated in Petri dishes containing PIM supplemented with 10 µL/L Tween 80 (Thermo Fisher Scientific, 28329) to minimize cell lysis^11^. Cultures were maintained in the dark at 22°C. In this study, PIM was prepared using an ammonium nitrate (NH₄NO₃)-free MS basal salt mixture (Ducefa Biochemie, M0238.0050), as elevated NH₄NO₃ concentrations are known to induce osmotic stress and cytotoxicity in protoplasts (Qiao et al. 1998). Liquid protoplast cultures were maintained in PIM for 7d, followed by transfer to liquid CIM1 for another 7d. Protoplasts were then collected by low-speed centrifugation, resuspended in a reduced volume of CIM1, and mixed with an equal volume of 3% sodium alginate solution. The alginate-protoplast suspension was poured onto a solid calcium agar bed and allowed to polymerize for 30 min. Emerging microcalli within the alginate layer were manually retrieved using fine forceps and transferred onto solid SIM medium for 28d to induce shoot regeneration. For standard or shortened alginate-based protocols, protoplasts were embedded in 3% alginate immediately after transfection using an equal volume of PIM. The cultures were maintained sequentially in PIM, CIM1, and CIM2 (Supplementary Table S3), with media changed every 5∼7d. Visible microcalli were transferred onto solid SIM for shoot induction. Regenerated shoots were excised and moved onto solid RIM for 21d to induce rooting. Fully regenerated plantlets were transferred into soil and acclimated in the growth chamber. Seeds from regenerated plants were harvested and stored at 4°C. The standard and shortened alginate culture protocols, PIM, CIM1, CIM2, and SIM-incubation periods were mentioned in each experimental setup. Protoplast cultures for Arabidopsis, *B. oleracea*, and *C. annuum* were maintained at 22°C, except for rice, tomato, and soybean, which were cultured at 25°C. Rice and soybean cultures used species-specific media compositions and timelines, as detailed in the corresponding experimental setups, figure legends, and Supplementary Table S3.

### Microscopy

Fluorescence microscopy (Carl Zeiss, AxioZoom V16) was performed to study protoplast viability, transfection efficiency, and microcalli formation in PIM and CIM. FDA-stained protoplasts were visualized using a GFP filter (excitation-emission at 488 nm). Transfected protoplasts expressing fluorescent proteins were imaged using three-channel fluorescence detection (green, red, and blue) with appropriate excitation lines and emission filters. CW-stained dividing protoplasts and developing microcalli were visualized using a DAPI filter (excitation-emission at 447 nm). Confocal microscopy (LSM700, Carl Zeiss) was used to detect nuclear-localized nYFP and SCR-mGFP signals in protoplasts and alginate-embedded microcalli transfected with tag-free SEPW constructs. Samples were sectioned, mounted on glass slides, and imaged using solid-state lasers (405-639 nm) and standard 3-channel settings alongside bright-field imaging. For root tip imaging, Arabidopsis seedlings homozygous for *pWOX5::nYFP* or *pSCR::SCR-mGFP* were stained with PI, washed with sterile water, and mounted on slides for confocal imaging. Images were acquired at 20X or 40X magnification with a frame rate of 5 fps. Image acquisition parameters (magnification, exposure, and gamma) were kept consistent across samples. Brightness and contrast adjustments were applied uniformly per channel using ZEN imaging software (Carl Zeiss).

### Protoplast quantification and microcalli assessment

Protoplast enumeration was conducted after isolation by using a hemacytometer under the light microscope (Carl Zeiss, Axioskop 40), as well as after transfection and staining experiments. ImageJ software was used to quantify the total number of protoplasts and microcalli. The percentage of microcalli formation was calculated as the ratio of microcalli to the total number of viable protoplasts. Additionally, ImageJ was also used to assess transfection efficiency by counting positive fluorescent cells (cells exhibiting fluorescence above the background threshold), measuring RFU to quantify fluorescence intensity (representing the signal from transfected cells relative to background fluorescence), and determining the percentage area of microcalli (microcalli area as a fraction of the field of view). Images were converted to 8-bit grayscale, followed by background subtraction to remove noise. Thresholding was applied using the Otsu method, with manual adjustments made to optimize contrast between microcalli and background. Particle analysis was performed with size criteria set between 50∼500 μm² and circularity between 0.3 and 1.0 to accurately detect microcalli while excluding particles touching the image edges. The ‘Analyze Particles’ function was used to count microcalli and measure their area, and manual validation was carried out to ensure the accuracy of detection. These settings ensured reliable identification and quantification of microcalli.

### Callus sectioning and staining

Microcalli from SIM were carefully picked using sterile forceps and embedded in a pre-prepared 2% (w/v) agarose block. Sectioning was performed using a vibratome (Leica, VT1200S) to obtain 100 µm-thick slices, which were subsequently collected in 1X PBS buffer. The sections were stained with 1% (w/v) toluidine blue O (Sigma-Aldrich, T3260) (Klimek-Chodacka et al. 2020). After staining, the sections were washed with 1X PBS buffer and visualized under a stereo microscope (Leica, Ivesta-3) to assess cellular structures.

### Genotyping

For the genotyping experiment, genomic DNA (gDNA) isolation from both Arabidopsis and crop plants was carried out following a well-established protocol (Edwards et al. 1991). The Edwards solution was diluted tenfold with TB buffer to create an extraction buffer. Approximately 5∼10 mg of leaf tissue was placed in an Eppendorf tube (e-tube) and crushed in 200 µL of the extraction buffer. The resulting green supernatant was carefully transferred to a fresh e-tube after centrifugation at 13,000 rpm for 5 min. Subsequently, gDNA was precipitated by adding an equal volume of isopropanol and washed with 70% ethanol. The precipitated gDNA was ultimately suspended in sterile water and utilized for PCR after quantification and quality assessment in an agarose gel. The primer sets employed for genotyping are detailed in Supplementary Table S2. PCR products were loaded and examined in an agarose gel to confirm the presence of gene-of-interest in the genome of regenerated plants. Finally, gel images were captured using the Gel Logic 100 imaging system.

### Generation of *pSCR::SCR-mGFP* and *pWOX5::nYFP* reporter lines

A 2,131 bp genomic fragment containing the *SCR* promoter and full-length coding region was amplified from Col gDNA using primers pSCR::SCR_Entry_F and pSCR::SCR_Entry_R (Supplementary Table S2), and cloned into the pENTR™/SD/D-TOPO vector (Invitrogen). The final *pSCR::SCR-mGFP* construct was generated by LR recombination into a pEarleyGate destination vector (Invitrogen, 11791020) and introduced into Col plants *via Agrobacterium tumefaciens*-mediated transformation using the floral dip method (Clough et al. 1998). Transgenic seedlings were selected on MS medium supplemented with 50 mg/L glufosinate ammonium (BASTA; GoldBio, P-165-1), and homozygous lines were established in the T3 generation. The *pWOX5::nYFP* reporter line (Shimotohno et al. 2018) was obtained from Ben Scheres (Wageningen University).

### Immunostaining

Nuclei were isolated from Arabidopsis and rice mesophyll protoplasts, either untransfected or transfected with AtSEPW and OsSEPW factors, respectively, at 3d post-transfection, and immunostaining was performed as described previously (Yelagandula et al. 2014). Primary antibodies against H3K4me3 (Abcam, ab8580) and H3K27me3 (Millipore, 07-449) were used at a 1:200 dilution. An Alexa Fluor 488-conjugated goat anti-rabbit secondary antibody (Invitrogen, A-11008) was used at a 1:500 dilution. Images were acquired using Airyscan 2 laser scanning confocal microscope (Carl Zeiss, LSM980) in super-resolution mode. Fluorescence signal intensities were quantified using the Histo function in ZEN software (Carl Zeiss).

### Single-cell RNA sequencing and data analysis

Arabidopsis mesophyll protoplasts were isolated from the leaves of 15-d-old seedlings as described previously (Yoo et al. 2007). The protoplasts were transfected with or without (mock) SEPW factors using 20% PEG and cultured for 2d in liquid PIM at 22°C in the dark. The mock samples were treated with only 20% PEG. The protoplasts were then filtered using a 40 μm cell strainer (SPL Life Sciences, 93040) to obtain a single-cell suspension culture. The protoplasts without any aggregates were then assayed for their viability using FDA staining (> 80% viable). Finally, the protoplasts were resuspended in 1X PBS buffer and counted using a haemocytometer before loading on a 10X Genomics single-cell instrument (Chromium iX, AS202115200) that generates single-cell Gel bead-in EMulsion (GEMs). The GEM preparation and barcoding were performed using the Chromium GEM-X Single Cell 3’ Chip Kit v4 (10X Genomics, PN-1000690). Post-GEM-RT cleanup and cDNA amplification were performed using the Chromium GEM-X Single Cell 3’ Kit v4 (10X Genomics, PN-1000691). The scRNA-seq libraries were constructed using the library construction kit C (10X Genomics, PN-1000694) and were sequenced with the Illumina sequencing platform NovaSeq 6000, by Macrogen Co., Inc (South Korea). The scRNA-seq sequencing reads were processed using Cell Ranger(v8.0.1;https://support.10xgenomics.com/single-cell-geneexpression/software/pipelines/latest/what-is-cell-ranger). TAIR10 reference genome and annotation files were downloaded from the Arabidopsis Information Resource (www.arabidopsis.org/). On average, 16,658 and 18,605 genes were detected in the mock and SEPW samples, respectively. The median unique molecular identifier (UMI) counts per cell on average were 455 for the mock and 908 for the SEPW samples. Pre-processing and quality control of scRNA-seq data were conducted using Seurat (v5) (Butler et al. 2014). The gene-cell matrices for each sample were imported into Seurat for downstream analysis. Low-quality cells containing more than 5% mitochondrial or chloroplast-derived reads were filtered out. The data were normalized using the “LogNormalize” method with a scale factor of 10,000. For dimensionality reduction, the “RunPCA” function was employed, followed by clustering using the “FindClusters” function at a resolution of 0.8. Cell embeddings were visualized in the Uniform Manifold Approximation and Projection (UMAP) space using the “RunUMAP” function. To assess the cell cycle stage of each cell, we utilized the “CellCycleScoring” function in Seurat, based on G1, S, and G2/M phase marker genes. Differential gene expression (DGE) analysis between mock and SEPW samples was performed using “FindMarkers” with the Wilcoxon rank sum test. Genes with an adjusted *p*-value < 0.05 were considered significant. Gene Ontology (GO) enrichment analysis of differentially expressed genes was conducted using the clusterProfiler (v4.0.5) package in R. Marker gene mapping and annotation strategies were adapted from a previously published study (Xu et al. 2021).

### Click-iT EdU imaging

Arabidopsis mesophyll protoplasts were transfected with SEPW (tag-free) constructs and cultured in liquid PIM for 2d prior to analysis. EdU incorporation was detected using the Click-iT™ EdU Imaging Kit (Invitrogen, C10339) according to the manufacturer’s instructions with minor modifications. Briefly, following incubation with 10μM EdU, protoplasts were harvested by gentle centrifugation and fixed in 4% paraformaldehyde (CellPick, P001-1000). To maintain osmotic stability during washes and permeabilization, buffers were prepared in 1X PBS supplemented with 0.5 M mannitol (PBS-M). Cells were permeabilized with 0.5% Triton X-100 in PBS-M and washed with 1% BSA (Fraction V) prepared in PBS-M prior to Click-iT reaction. After fluorescent azide labeling, nuclei were counterstained with Hoechst 33342 (Invitrogen, C10337) to visualize nuclear DNA. Samples were mounted on poly-L-lysine coated slides and imaged by confocal microscopy (Carl Zeiss, LSM700) under identical acquisition settings across conditions.

### RT-qPCR analysis

Soybean mesophyll protoplasts were transfected with or without (mock control) GmSEPW and cultured in liquid PIM for 2d. Cells were harvested by low-speed centrifugation (600 rpm, 5 min, 4°C), and total RNA was extracted using the RNeasy Plant Mini Kit (Qiagen, 74904) according to the manufacturer’s instructions. RNA integrity was assessed by 2% agarose gel electrophoresis, and quantified using the NanoDrop (GE Healthcare). 2 μg of high-quality total RNA was used for first-strand cDNA synthesis using the RevertAid First Strand cDNA Synthesis Kit (Thermo Fisher Scientific, K1621). Following reverse transcription (RT), quantitative PCR (qPCR) was performed on first-strand DNA with a real-time PCR cycler (QIAGEN Rotor-Gene Q 2Plex) by using TOPreal SYBR Green qPCR PreMIX (Enzynomics, RT500M). Gene expression was quantified using standard curve-based calculations, and transcript levels were normalized to the internal control gene *GmActin4*. Primers used for RT-qPCR analysis are listed in Supplementary Table S2.

### Statistical analysis

All data are presented as mean ± s.d., based on a minimum of two independent biological replicates, each comprising three technical replicates unless otherwise stated. Statistical analyses were performed using one-way or two-way ANOVA, as appropriate, with significance determined at *P* ≤ 0.05. Analyses were conducted using GraphPad Prism 8.0. ScRNA-seq experiments were carried out in two independent biological replicates. Downstream statistical analyses and data visualizations for scRNA-seq were performed using R (v4.2.0). Additional details, including replicate numbers, statistical tests, and exact *P* values, are provided in the figure legends.

## Acknowledgements

We thank Suk Weon Kim (KRIBB), Suk-Yoon Kwon (KRIBB), Byoung-Cheorl Kang (SNU), Do-Soon Kim (SNU), Ki-Hong Jung (KU), and the National Agrobiodiversity Center of Korea (https://genebank.rda.go.kr) for crop seeds used in this study. *pSCR::SCR-mGFP* transgenic plant was generated by Myung-Hwan Choi (SNU).

## Author contributions

Y.S.N conceived the idea, designed, and supervised the research. G.S.J performed the mainstream experiments including vector construction, Arabidopsis gene cloning, establishing transfection and microscopic analyses. G.S.J and M.N.K shared the work related to Arabidopsis regeneration. M.N.K performed the work related to cabbage, tomato, and hot pepper regeneration with G.S.J’s co-contribution to the work of hot pepper. Rice gene cloning and regeneration work were performed by G.S.J, while soybean gene cloning, expression analysis, and regeneration work were performed by M.N.K and B.N. Experiments related to scRNA-seq and EdU assay were performed by G.S.J, while the immunostaining of Arabidopsis and rice nuclei was performed by C.P. G.S.J and Y.S.N analyzed data. G.S.J and Y.S.N made figures and wrote the manuscript. All authors read and approved the manuscript.

## Data availability

The data supporting the findings of this study are available in the article and the supplementary information. The full-length sequences of the HBT-PGSG-sGFP/mRFP/mBFP-NOS vectors have been deposited in GenBank under the accession numbers PP959378∼PP959380. The scRNA-seq data have been deposited in the Gene Expression Omnibus under the SuperSeries accession number GSE307300.

## Supplementary data

The online version contains supplementary material available at https://.

